# RNA-binding protein Syncrip regulates Starvation-Induced Hyperactivity in adult *Drosophila*

**DOI:** 10.1101/2020.01.07.897652

**Authors:** Wanhao Chi, Wei Liu, Wenqin Fu, Shengqian Xia, Ellie S. Heckscher, Xiaoxi Zhuang

## Abstract

How to respond to starvation determines fitness. One prominent behavioral response is increased locomotor activities upon starvation, also known as Starvation-Induced Hyperactivity (SIH). SIH is paradoxical as it promotes food seeking but also increases energy expenditure. Despite its importance in fitness, the genetic contributions to SIH as a behavioral trait remains unexplored. Here, we examined SIH in the *Drosophila melanogaster* Genetic Reference Panel (DGRP) and performed genome-wide association studies. We identified 23 significant loci, corresponding to 14 genes, significantly associated with SIH in adult *Drosophila*. Gene enrichment analyses indicated that genes encoding ion channels and mRNA binding proteins (RBPs) were most enriched in SIH. We are especially interested in RBPs because they provide a potential mechanism to quickly change protein expression in response to environmental challenges. Using RNA interference, we validated the role of *syp* in regulating SIH. *syp* encodes Syncrip (Syp), an RBP. While ubiquitous knockdown of *syp* led to semi-lethality in adult flies, adult flies with neuron-specific *syp* knockdown were viable and exhibited decreased SIH. Using the Temporal and Regional Gene Expression Targeting (TARGET) system, we further confirmed the role of Syp in adult neurons in regulating SIH. To determine how *syp* is regulated by starvation, we performed RNA-seq using the heads of flies maintained under either food or starvation conditions. RNA-seq analyses revealed that *syp* was alternatively spliced under starvation while its expression level was unchanged. We further generated an alternatively-spliced-exon-specific knockout (KO) line and found that KO flies showed reduced SIH. Together, this study demonstrates a significant genetic contribution to SIH as a behavioral trait, identifies *syp* as a SIH gene, and highlights the significance of RBPs and post-transcriptional processes in the brain in regulating behavioral responses to starvation.

**Author summary:** Animals living in the wild often face periods of starvation. How to physiologically and behaviorally respond to starvation is essential for survival. One behavioral response is Starvation-Induced Hyperactivity (SIH). We used the *Drosophila melanogaster* Genetic Reference Panel, derived from a wild population, to study the genetic basis of SIH. Our results show that there is a significant genetic contribution to SIH in this population, and that genes encoding RNA binding proteins (RBPs) are especially important. Using RNA interference and the TARGET system, we confirmed the role of an RBP Syp in adult neurons in SIH. Using RNA-seq and Western blotting, we found that *syp* was alternatively spliced under starvation while its expression level was unchanged. Further studies from *syp* exon-specific knockout flies showed that alternative splicing involving two exons in *syp* was important for SIH. Together, this study identifies *syp* as a SIH gene and highlights an essential role of post-transcriptional modification in regulating this behavior.

## Introduction

Animals living in the natural environment often experience periods of starvation, and they have thus developed different physiological and behavioral strategies to respond to starvation [1, 2]. One well-documented behavioral response is Starvation-Induced Hyperactivity (SIH), that is, animals will increase their locomotor activity upon starvation [3]. SIH has been observed in both flies [4–9] and mammals [10–12], suggesting that this behavior is evolutionarily conserved. From the viewpoint of energy gain and expenditure, SIH seems paradoxical. On one hand, it facilitates food acquisition and energy intake when food is available [7]; on the other hand, it increases energy expenditure and makes starved animals even more vulnerable when food is not available [4]. Therefore, genetic dispositions to either too much or too little SIH would impair fitness depending on the environment. However, the genetic contributions to SIH as a behavioral trait remains unexplored.

Recent studies suggest that SIH is highly regulated. A number of genes have been shown to regulate SIH in adult *Drosophila*, including genes encoding energy sensors [5, 6], neuropeptides/neuropeptide receptors [4, 8], and neurotransmitters [7]. Recently, *dG9a* was shown to regulate SIH in adult *Drosophila* [9]. *Gene dG9a* encodes a histone methyltransferase, suggesting that SIH is also regulated at the epigenetic level. In addition, post-transcriptional modifications, especially alternative mRNA splicing, has been shown to be effective for cells and animals to quickly respond to starvation as well as other stresses [13–15]. However, whether or not alternative mRNA splicing plays a role in SIH has not been reported.

In this study, we first set out to determine whether SIH varied in a population of *Drosophila melanogaster* with a diverse genetic background. We used the *Drosophila melanogaster* Genetic Reference Panel (DGRP). The DGRP consists of 205 inbred wild-type strains derived from a single population that was collected from Raleigh, North Carolina, USA [16, 17]. It is a recently established community resource and has been used to examine the genetic basis of more than 60 quantitative traits [18]. We quantified SIH in DGRP and confirmed a significant genetic contribution to SIH. We then performed genome-wide association studies and identified 23 significant loci from 14 genes associated with SIH. We found that genes with ion channel activities and mRNA binding activities were especially enriched in SIH. Using RNA interference, we validated the role of a gene encoding an RNA-binding protein Syncrip (Syp) [19] in neurons in regulating SIH. Using the Temporal and Regional Gene expression Targeting (TARGET) system [20], we further confirmed the role of Syp in adult neurons in regulating SIH. To determine how *syp* was regulated by starvation, we performed RNA-seq on the heads of flies maintained under either food or starvation conditions. We found that *syp* was alternatively spliced under starvation while its expression level remained unchanged. We further generated an alternatively-spliced-exon-specific knockout (KO) line. Activity data from KO flies showed that alternative splicing involving two exons in *syp* was important for SIH.

## Results

### Natural variation in Starvation-Induced Hyperactivity across DGRP

To study the natural variation in Starvation-Induced Hyperactivity (SIH), we monitored the locomotor activity of 198 strains in the *Drosophila melanogaster* Genetic Reference Panel (DGRP) [16, 17] under either food or starvation conditions in the Drosophila Activity Monitor for three days (S1 Fig; See Methods). DGRP consists of 205 inbred strains. We excluded those that did not breed well.

The locomotor activity of each strain under either food or starvation conditions was plotted as a function of time, sixty hours in total. Representative plots are shown in Fig 1A. We observed that, under the 12-hr:12-hr day-night condition and with the presence of food, all strains showed typical bimodal activity peaks, a day-night rhythm governed by the circadian system [21] (black lines in Fig 1A). Such rhythmic activities were disrupted by food deprivation in most strains. Starved flies became persistently active and the hyperactivity could occur in either day or night (red lines in Fig 1A). There were exceptions as some strains did not show obvious SIH (See a representative strain in Fig 1A, the bottom panel).

**Fig 1.**
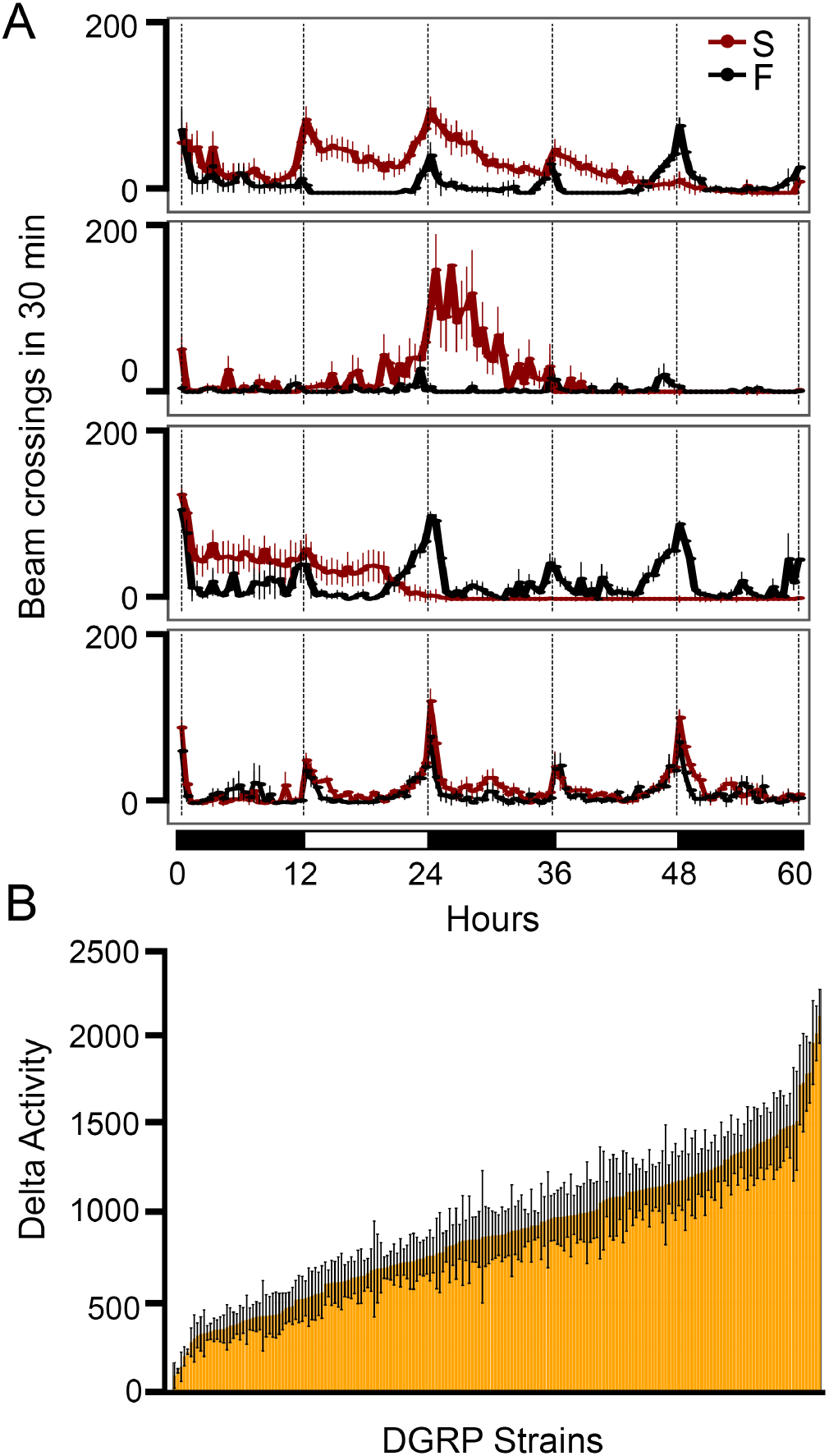
Natural variation of Starvation-Induced Hyperactivity in the DGRP. (A) Representative plots of locomotor activity responses to starvation. Beam crosses in 30 min were plotted as a function of time. Black lines represent activities under the food condition (F) and red lines represent activities under the starvation condition (S). Black and white bars at the bottom of panel A represent night and day cycles, respectively. Each plot was from one DGRP strain (n = 8 per condition). (B) Natural variation occurred in SIH across DGRP. Error bars represent SEM.

To quantify SIH, we used the total activity during the 12-hr nighttime or daytime, and then calculated the activity difference (Delta Activity, DA) between the starvation condition and the food condition (See Methods for details). Since activities were recorded for a total of sixty hours (3 nights and 2 days), a total of five DA values were obtained accordingly. To compare the DAs among 198 trains in the DGRP, we used the largest DA among five DAs. DA is the absolute activity difference from baseline. We did not use the activity ratio over baseline because we did not see a correlation between DA and baseline activity (*ρ* = 0.046, *P* = 0.519; S1 Table), suggesting that DA was not increased proportionally to the baseline level. A summary plot of the largest DA (DA hereafter) across 198 strains was shown in Fig 1B. DA ranged from 93.61 ±72.22 to 2101.91 ±150.76 with a broad sense heritability of *H*^2^ = 0.38 (S1 Table, S2 Table). Thus, SIH varies among DGRP strains and it has a strong genetic basis.

### Starvation-Induced Hyperactivity is negatively correlated with Starvation Resistance

Previous studies showed that flies with reduced SIH survived longer under starvation [5, 6], suggesting that SIH is negatively correlated with Starvation Resistance (SR). To examine whether this relationship could also be observed in DGRP strains which were derived from a natural population, we did a correlation analysis between SIH and SR. We found that SIH was indeed negatively correlated with SR, but the correlation was not particularly strong (*ρ* = −0.231, *P* = 1.04e-03; n =198). Studies have demonstrated that SIH resembles foraging behavior [7]. Previously, we have studied the survival rates of DGRP strains in a foraging environment [22]. Thus, we further examined the correlation relationship between SIH and foraging survival rates of DGRP strains. We did not find any correlation between them (*ρ* = −0.073, *P* = 0.309; n = 197), which is not too surprising as the beneficial or detrimental effects of SIH may well depend on the specific environmental conditions.

### Genome-wide association analysis of Starvation-Induced Hyperactivity

We next performed genome-wide association (GWA) analyses using the DGRP analysis pipeline (http://dgrp2.gnets.ncsu.edu) [16, 17]. In total, we identified 23 SNPs/Indels that were associated significantly with SIH (*P* < 1 × 10^−5^, Fig 2A, Table 1, S2 Fig, S3 Table). Among all these significant loci, 43.5% of them are located in the intronic region, 43.5% are in the intergenic region or less than 1 kb downstream or upstream of an annotated gene and 13.0% are in the coding region (Fig 2B), which is comparable to the distribution of SNPs/Indels from previous studies in the DGRP (*P* = 0.4428, Fisher’s exact test) [23].

**Table 1.**
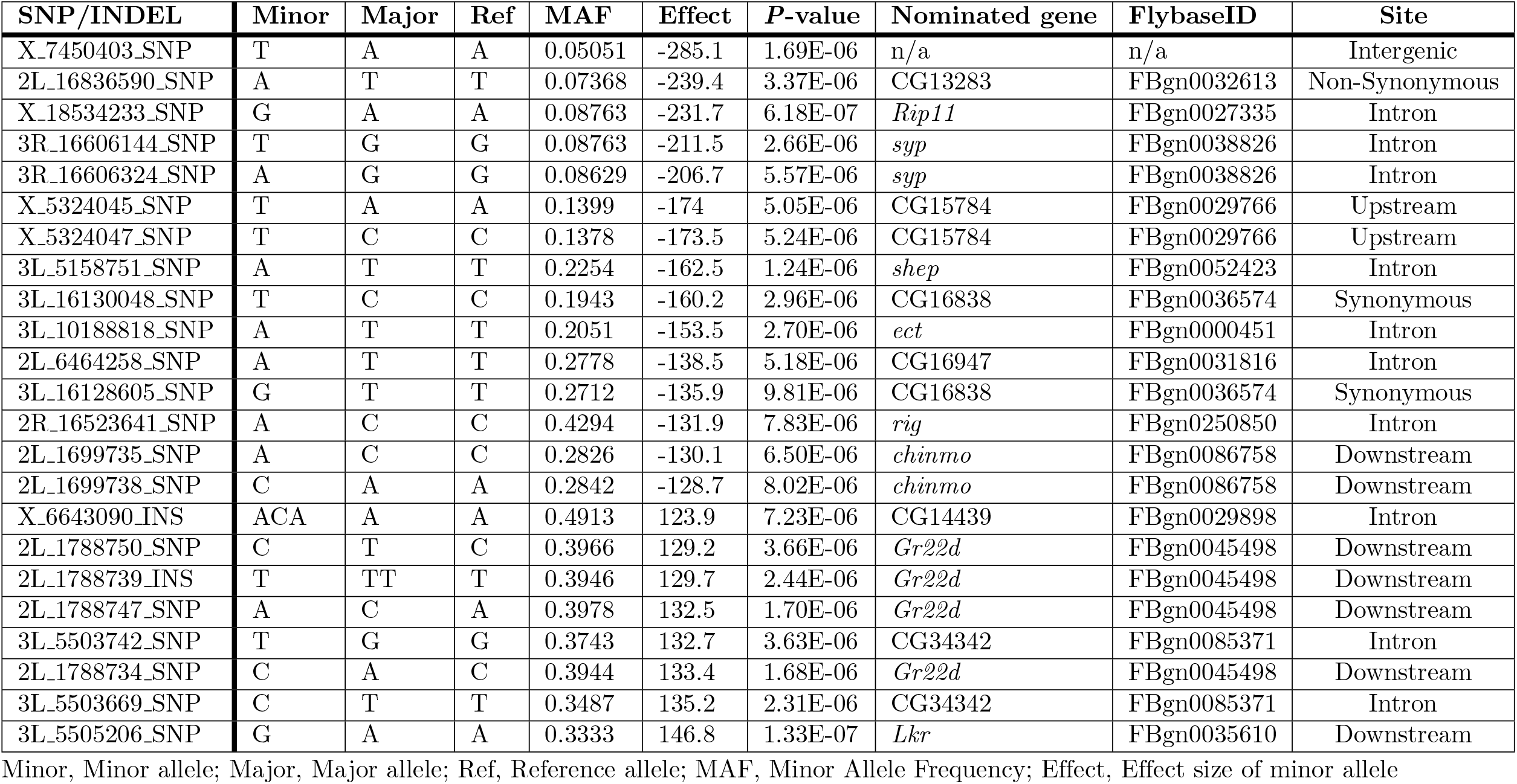
SNPs/Indels significantly associated with SIH variation at *P* < 1 × 10^−5^.

| SNP/INDEL | Minor | Major | Ref | MAF | Effect | P-value | Nominated gene | FlybaseID | Site |
| --- | --- | --- | --- | --- | --- | --- | --- | --- | --- |
| X_7450403_SNP | T | A | A | 0.05051 | -285.1 | 1.69E-06 | n/a | n/a | Intergenic |
| 2L_16836590_SNP | A | T | T | 0.07368 | -239.4 | 3.37E-06 | CG13283 | FBgn0032613 | Non-Synonymous |
| X_18534233_SNP | G | A | A | 0.08763 | -231.7 | 6.18E-07 | *Rip11* | FBgn0027335 | Intron |
| 3R_16606144_SNP | T | G | G | 0.08763 | -211.5 | 2.66E-06 | *syp* | FBgn0038826 | Intron |
| 3R_16606324_SNP | A | G | G | 0.08629 | -206.7 | 5.57E-06 | *syp* | FBgn0038826 | Intron |
| X_5324045_SNP | T | A | A | 0.1399 | -174 | 5.05E-06 | CG15784 | FBgn0029766 | Upstream |
| X_5324047_SNP | T | C | C | 0.1378 | -173.5 | 5.24E-06 | CG15784 | FBgn0029766 | Upstream |
| 3L_5158751_SNP | A | T | T | 0.2254 | -162.5 | 1.24E-06 | *shep* | FBgn0052423 | Intron |
| 3L_16130048_SNP | T | C | C | 0.1943 | -160.2 | 2.96E-06 | CG16838 | FBgn0036574 | Synonymous |
| 3L_10188818_SNP | A | T | T | 0.2051 | -153.5 | 2.70E-06 | *ect* | FBgn0000451 | Intron |
| 2L_6464258_SNP | A | T | T | 0.2778 | -138.5 | 5.18E-06 | CG16947 | FBgn0031816 | Intron |
| 3L_16128605_SNP | G | T | T | 0.2712 | -135.9 | 9.81E-06 | CG16838 | FBgn0036574 | Synonymous |
| 2R_16523641_SNP | A | C | C | 0.4294 | -131.9 | 7.83E-06 | *rig* | FBgn0250850 | Intron |
| 2L_1699735_SNP | A | C | C | 0.2826 | -130.1 | 6.50E-06 | *chinmo* | FBgn0086758 | Downstream |
| 2L_1699738_SNP | C | A | A | 0.2842 | -128.7 | 8.02E-06 | *chinmo* | FBgn0086758 | Downstream |
| X_6643090_INS | ACA | A | A | 0.4913 | 123.9 | 7.23E-06 | CG14439 | FBgn0029898 | Intron |
| 2L_1788750_SNP | C | T | C | 0.3966 | 129.2 | 3.66E-06 | *Gr22d* | FBgn0045498 | Downstream |
| 2L_1788739_INS | T | TT | T | 0.3946 | 129.7 | 2.44E-06 | *Gr22d* | FBgn0045498 | Downstream |
| 2L_1788747_SNP | A | C | A | 0.3978 | 132.5 | 1.70E-06 | *Gr22d* | FBgn0045498 | Downstream |
| 3L_5503742_SNP | T | G | G | 0.3743 | 132.7 | 3.63E-06 | CG34342 | FBgn0085371 | Intron |
| 2L_1788734_SNP | C | A | C | 0.3944 | 133.4 | 1.68E-06 | *Gr22d* | FBgn0045498 | Downstream |
| 3L_5503669_SNP | C | T | T | 0.3487 | 135.2 | 2.31E-06 | CG34342 | FBgn0085371 | Intron |
| 3L_5505206_SNP | G | A | A | 0.3333 | 146.8 | 1.33E-07 | *Lkr* | FBgn0035610 | Downstream |

Minor, Minor allele; Major, Major allele; Ref, Reference allele; MAF, Minor Allele Frequency; Effect, Effect size of minor allele

**Fig 2.**
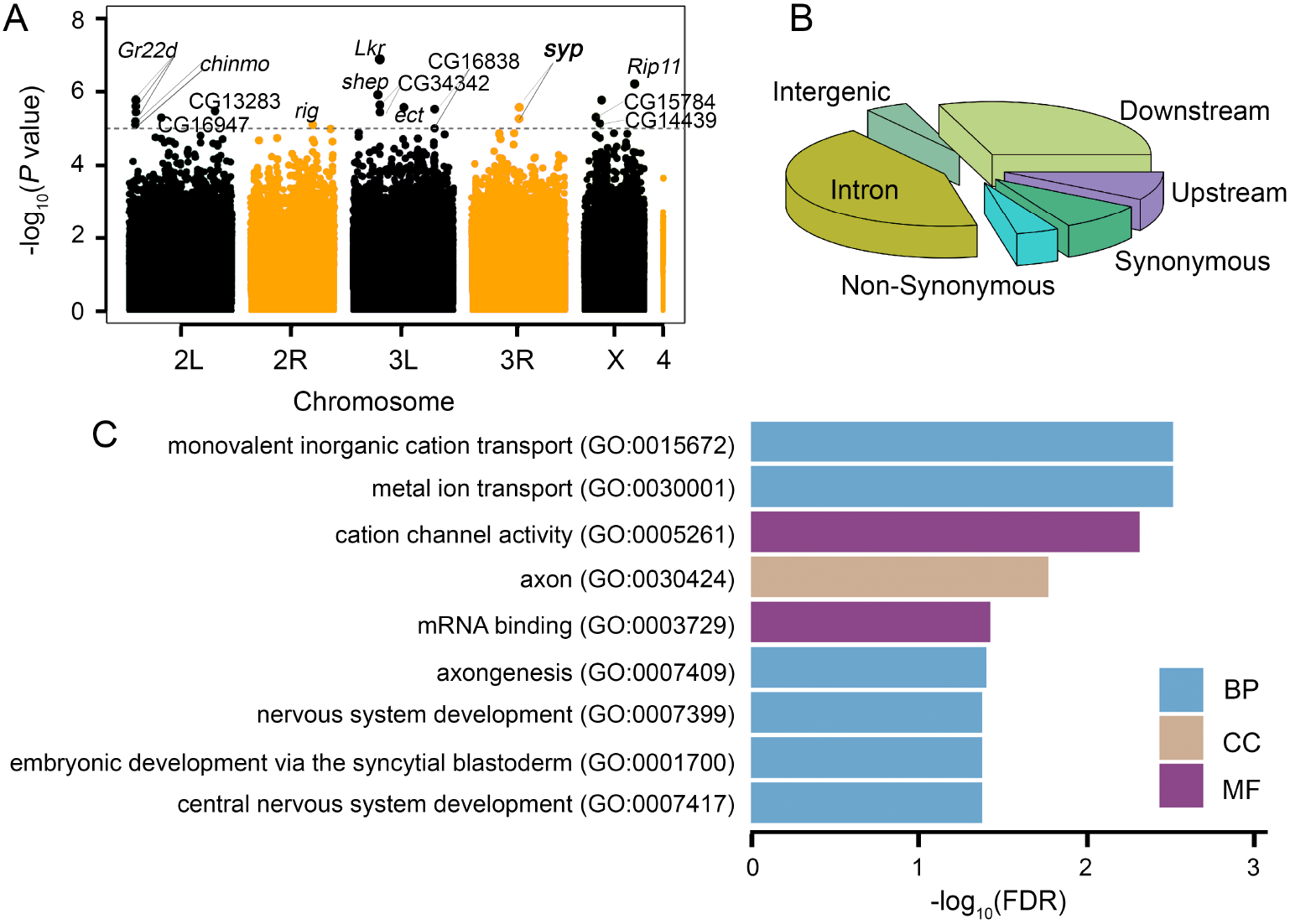
Manhattan plot for GWA, genomic location of significant SNPs, and GO analysis. (A) Manhattan plot of SIH. The gray dotted line is the significant cutoff line used for gene nomination (*P* < 1 × 10^−5^). (B) Genomic location of significant SNPs/Indels. (C) GO terms associated with more than four genes. BP: Biological Process; CC: Cellular Component; MF: Molecular Function.

### Genes associated with SIH are enriched in molecular function GO terms related to ion channels and mRNA binding

A total of 14 genes were nominated (Table 2). To study if a particular molecular function was enriched in SIH, we performed gene enrichment analysis [24, 25]. Since 14 candidate genes with *P* < 1 × 10^−5^ in GWA were too few for gene enrichment analyses, we therefore relaxed the cutoff threshold and did a step-wise relaxation using different cutoffs in the range of *P* < 1 × 10^−5^ and *P* < 1 × 10^−4^. The number of candidate genes associated with *P* < 2 × 10^−5^, *P* < 4 × 10^−5^, *P* < 6 × 10^−5^, *P* < 8 × 10^−5^, and *P* < 1 × 10^−4^ is 27, 42, 63, 89, and 112, respectively (S4 Table). We performed Gene Ontology (GO) analyses for candidate genes from the cutoffs of *P* < 1 × 10^−4^ and *P* < 8 × 10^−5^ since the other cutoffs yielded too few candidates for GO analyses. We found that candidate genes were enriched in 54 and 27 GO terms, respectively (S5 Table). Terms associated with more than four candidate genes with the cutoff of *P* < 1 × 10^−4^ were shown in Fig 2C. Among them, molecular function GO terms are cation channel activity (GO:0005261) and mRNA binding (GO:0003729). It is worth noting that mRNA binding (GO:0003729) was also associated with candidate genes with the cutoff of *P* < 8 × 10^−5^ (S5 Table), suggesting that genes with this molecular function are particularly important for SIH.

**Table 2.**
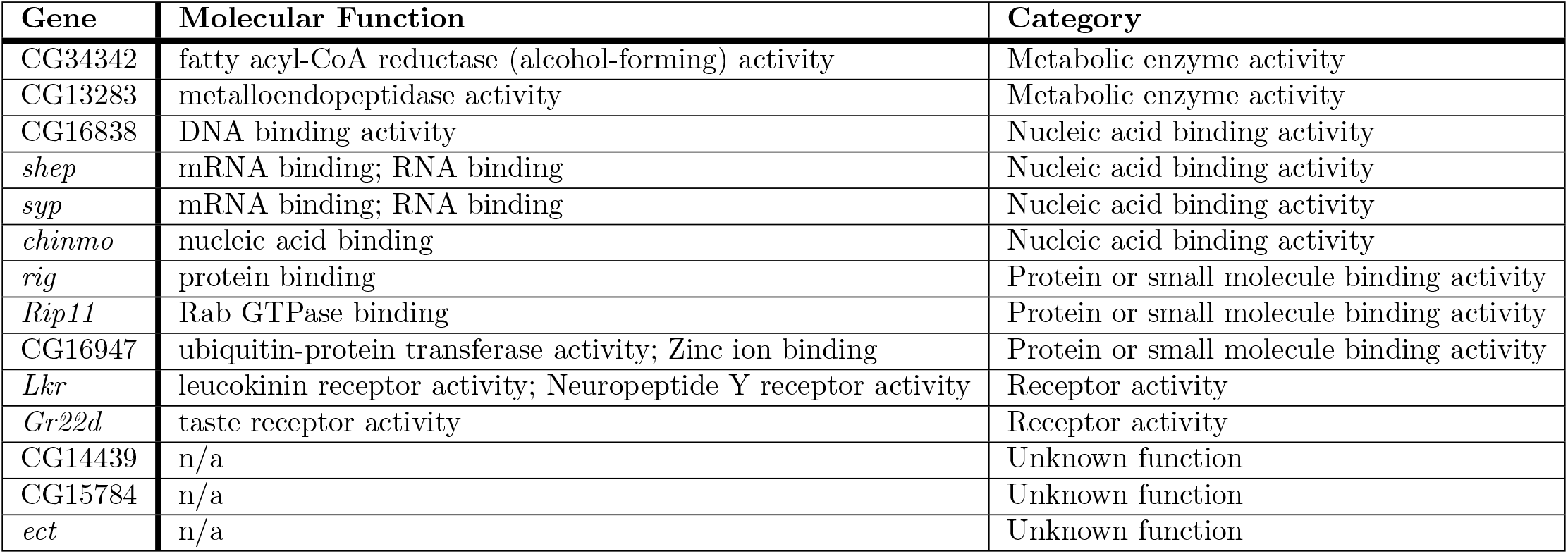
Known or predicted molecular functions of nominated genes.

| Gene | Molecular Function | Category |
| --- | --- | --- |
| CG34342 | fatty acyl-CoA reductase (alcohol-forming) activity | Metabolic enzyme activity |
| CG13283 | metalloendopeptidase activity | Metabolic enzyme activity |
| CG16838 | DNA binding activity | Nucleic acid binding activity |
| *shep* | mRNA binding; RNA binding | Nucleic acid binding activity |
| *syp* | mRNA binding; RNA binding | Nucleic acid binding activity |
| *chinmo* | nucleic acid binding | Nucleic acid binding activity |
| *rig* | protein binding | Protein or small molecule binding activity |
| *Rip11* | Rab GTPase binding | Protein or small molecule binding activity |
| CG16947 | ubiquitin-protein transferase activity; Zinc ion binding | Protein or small molecule binding activity |
| *Lkr* | leucokinin receptor activity; Neuropeptide Y receptor activity | Receptor activity |
| *Gr22d* | taste receptor activity | Receptor activity |
| CG14439 | n/a | Unknown function |
| CG15784 | n/a | Unknown function |
| *ect* | n/a | Unknown function |

In addition to GO analysis, we also grouped all significant candidate genes into five categories based on their known or predicted molecular functions from the fly database (Table 2). Category A includes genes encoding proteins with metabolic enzyme activities (2/14); Category B includes genes encoding proteins with nucleic acid (both DNA and RNA) binding activities (4/14); Category C includes genes encoding proteins with binding activities to proteins or small molecules (3/14); Category D includes genes encoding proteins with receptor activities (2/14). Lastly, Category E includes genes with still unknown molecular functions (3/14).

We were most intrigued by genes encoding proteins that have mRNA binding activities since many mRNA binding proteins (RBPs) are splicing regulators for alternative mRNA splicing, one of the important cellular responses that are often observed when cells/organisms are under starvation as well as other stresses [13–15]. Two genes that encode RNA binding proteins were shown in the list of nominated genes: *syp* (*syncrip*) and *shep* (*alan shepard*). We chose to validate gene *syp* as the loss of *shep*, even only in neurons, can lead to abnormal locomotor activities in adult flies [26], which could complicate the final data interpretation.

### Knockdown of *syp* in neurons significantly affects SIH

To test whether gene *syp* plays a role in SIH, we used the binary UAS/GAL4 system [27] to knock down its expression. We first crossed the gene-specific UAS-RNAi line with an *actin*-Gal4 driver to generate ubiquitous *syp* knockdown flies. We observed that ubiquitous *syp* knockdown led to lethality before the adult stage. This observation is consistent with previous studies showing that *syp* mutant flies have highly decreased adult viability [28]. Given that *syp* is highly expressed in adult heads (modENCODE.org), we thus generated neuron-specific *syp* knockdown flies by crossing UAS-*syp* RNAi flies with a pan-neuronal driver *nSyb*-Gal4. Neuron-specific *syp* knockdown flies survived well, suggesting that the fatality of ubiquitous *syp* knockdown flies is likely due to the lack of expression of *syp* in non-neuronal cells.

We then tested neuron-specific *syp* knockdown and control flies in the Drosophila Activity Monitor under either food or starvation conditions. As shown in Fig 3A, neuron-specific *syp* knockdown did not affect the locomotor activities in the food condition, suggesting that *syp* in neurons is not required for the general locomotor activity. However, Delta Activity from them was significantly lower than that from control flies (Fig 3B), demonstrating a specific role of *syp* in SIH. The reduced Delta Activity was also observed in neuron-specific *syp* knockdown flies generated from another independent UAS-*syp* RNAi line (Fig 3C and 3D). We therefore conclude that neuron-specific knockdown of *syp* impairs SIH.

**Fig 3.**
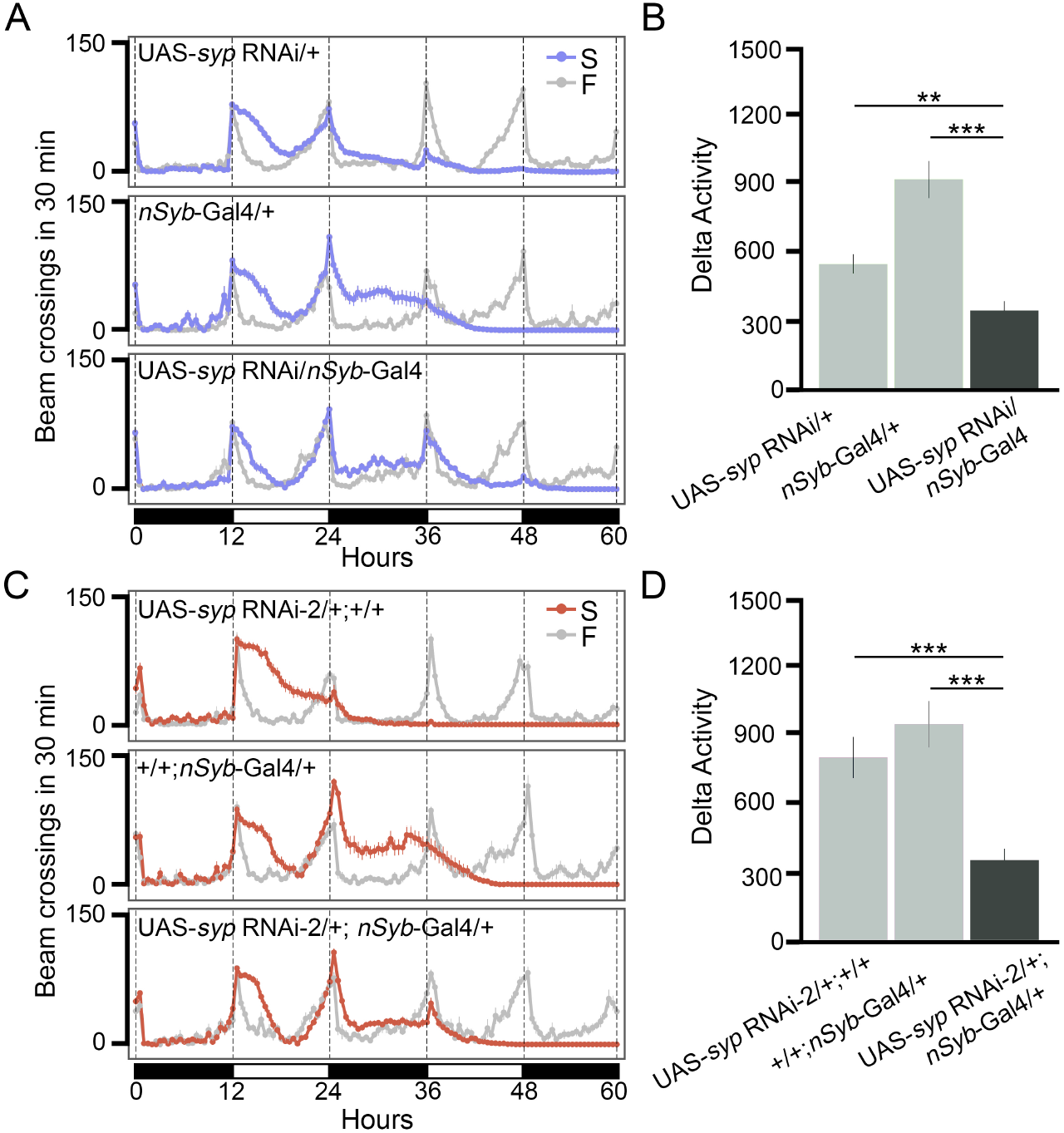
Knocking down of *syp* in neurons reduces SIH. (A) Locomotor activity responses to starvation from neuron-specific *syp* knockdown flies (genotype: UAS-*syp* RNAi/*nSyb*-Gal4) and two parental control lines (genotypes: UAS-*syp* RNAi/+ and *nSyb*-Gal4/+, respectively). Gray lines represent activities under the food condition (F) and purple lines represent activities under the starvation condition (S). Black and white bars at the bottom of the panel represent night and day cycles, respectively. n = 16-20 per genotype per condition. (B) Summary plot of SIH in panel A. (C) Locomotor activity responses to starvation from neuron-specific *syp* knockdown flies generated from a second RNAi line (genotype: UAS-*syp* RNAi-2/+; *nSyb*-Gal4/+) and two parental control lines (genotypes: UAS-*syp* RNAi-2/+; +/+ and +/+; *nSyb*-Gal4/+, respectively). n = 20-24 per genotype per condition. (D) Summary plot of SIH in panel C. ** *P* < 0.01, *** *P* < 0.001, unpaired *t*-test with Bonferroni Correction. Error bars represent SEM.

### Syp in adult neurons regulates Starvation-Induced Hyperactivity

Syp has been shown to affect synaptic morphology and vesicle release at the neuromuscular junction in *Drosophila* larvae [29, 30]. Moreover, it has also been implicated in neuron/glia cell fate determination during development [31, 32]. The role of *syp* in regulating SIH, therefore, could be due to its developmental effects. To rule out the possibility that Syp regulates SIH due to its developmental effects, we combined UAS-*syp* RNAi with the Temporal and Regional Gene Expression Targeting (TARGET) system [20] to control the expression of RNAi temporally. The TARGET system consists of three elements: Gal4, UAS, and temperature sensitive Gal80 mutant (Gal80^ts^). The Gal80^ts^ blocks Gal4-induced RNAi expression when tissue is exposed to the Gal80^ts^ permissive temperature (18°C). In contrast, at a Gal80^ts^ restrictive temperature (31°C), Gal80^ts^ loses its binding to Gal4, which allows Gal4-dependent RNAi expression. Therefore, breeding flies at 18°C and testing them at 31°C allowed us to bypass the developmental effects of *syp* KD.

We first bred flies at 18°C until adult flies emerged. We then shifted the temperature of the incubator to 31°C and incubated adult flies in the new food vials for one more day, which was followed by behavioral testing at 31°C under both starvation and food conditions for three days (Fig 4A). Results showed that *Tub*-Gal80^ts^/+; *nSyb*-Gal4/UAS-*syp* RNAi (ts-*syp*-KD) flies exhibited significantly attenuated SIH in comparison to control flies (genotypes: +/+; *nSyb*-Gal4/+, *Tub*-Gal80^ts^/+; UAS-*syp* RNAi/+, and *Tub*-Gal80^ts^/+; +/+. Fig 4B and 4C), and that the Delta Activity level from ts-*syp*-KD flies was comparable to that from +/+; *nSyb*-Gal4/UAS-*syp* RNAi (*syp*-KD) flies (*P* = 0.6863), demonstrating that knocking down of *syp* in the adult stage is sufficient for suppression of SIH.

**Fig 4.**
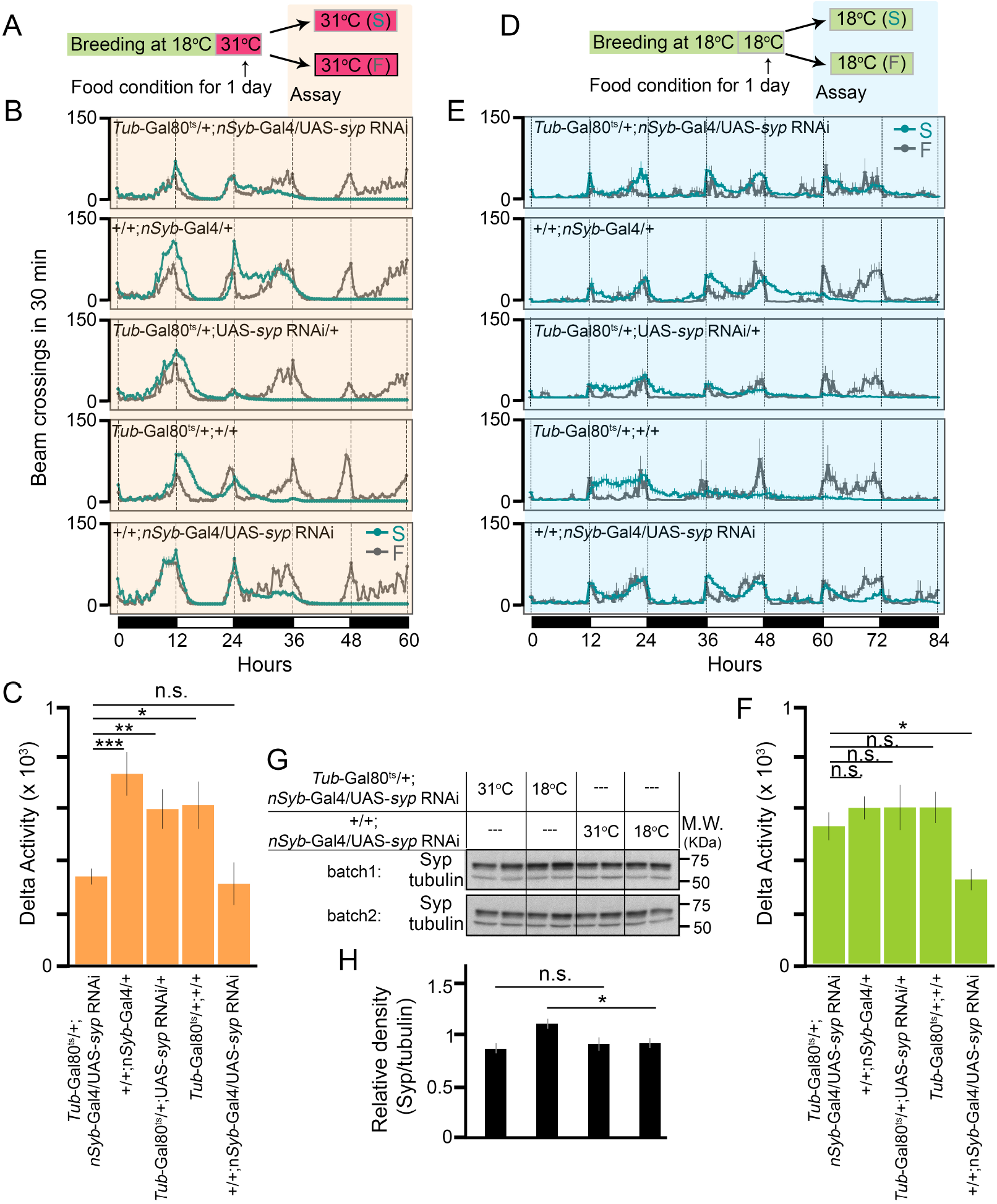
Syp in adult neurons regulates SIH. (A) Breeding and testing scheme. S: starvation condition; F: food condition. (B) Locomotor activity responses to starvation from each genotype under the condition shown in panel A. Gray lines represent activities under the food condition (F) and green lines represent activities under the starvation condition (S). n = 24-30 per genotype per condition. (C) Summary plot of SIH from each genotype in panel B. (D) Breeding and testing scheme. S: starvation condition; F: food condition. (E) Locomotor activity responses to starvation from each genotype under the condition shown in panel D. n = 8-28 per genotype per condition. (F) Summary plot of SIH from each genotype in panel E. (G) Western blot of adult fly head homogenate from *Tub*-Gal80^ts^/+;*nSyb*-Gal4/UAS-*syp* RNAi (ts-*syp*-KD) and +/+;*nSyb*-Gal4/UAS-*syp* RNAi (*syp*-KD) flies. Four biological replicates per genotype per condition. Tubulin is the loading control. (H) Quantification of Syp from four biological replicates. * *P* < 0.05, ** *P* < 0.01, *** *P* < 0.001, n.s.: *P* > 0.05, unpaired *t*-test with Bonferroni Correction. Error bars represent SEM.

To verify that Gal80^ts^ did inhibit Gal4 and prevent *syp* KD at 18°C, we also bred and tested adult flies all at 18°C (Fig 4D). Under this condition, as shown in Fig 4E and 4F, ts-*syp*-KD exhibited a similar level of SIH to control flies and a significant higher level of SIH than *syp*-KD flies. To further confirm that *syp* was indeed knocked down in adult flies at 31°C but not at 18°C, we compared the Syp protein levels in adult fly heads from ts-*syp*-KD and *syp*-KD flies that were maintained at either 31°C or 18°C for two days. As shown in Fig 4G and 4H, at 31°C, the Syp level was comparable between two genotypes (*P* = 0.4476). In comparison, at 18°C, the Syp level in ts-*syp*-KD flies was about 20% higher than that in *syp*-KD flies (*P* = 0.0496). The relatively moderate difference between them is most likely due to non-neurons mixed in the quantification of proteins in the fly head. Taken together, we conclude that it is Syp in adult neurons that regulates SIH.

### *syp* is alternatively spliced upon starvation

Gene *syp* spans a region of 54 kb in the genome, and it has 20 well-documented splice variants (www.flybase.org, Fig 5). To study how *syp* was regulated under starvation, we performed RNA sequencing (RNA-seq) on adult heads from control flies (*w*^1118^) reared under either food or starvation conditions (S3 Fig, S4 Fig, S5 Fig, S6 Fig; see Methods for details). We first examined whether the *syp* mRNA level was altered after starvation. Results from differential expression analyses showed that a total of 574 genes were down-regulated, and a total of 84 genes were up-regulated under starvation (Criteria applied: false-discovery rate (FDR) < 0.05 and fold change ≥2 or fold change ≤ −2; S6 Table). Gene *syp* was not on the list, suggesting that its expression was not dramatically regulated by starvation. This conclusion was further corroborated by similar protein levels of *syp* under two conditions (S7 Fig). To examine the functional implications of down-versus up-regulated genes, we performed gene enrichment analyses for each list of genes. We found that down-regulated genes participate in a variety of metabolic pathways, whereas up-regulated genes are mainly enriched in pathways related to DNA repair, nutrients recycling, and spliceosome (S6 Fig). The fact that genes involved in the spliceosome are upregulated upon starvation suggests that alternative splicing may underlie responses to starvation.

**Fig 5.**
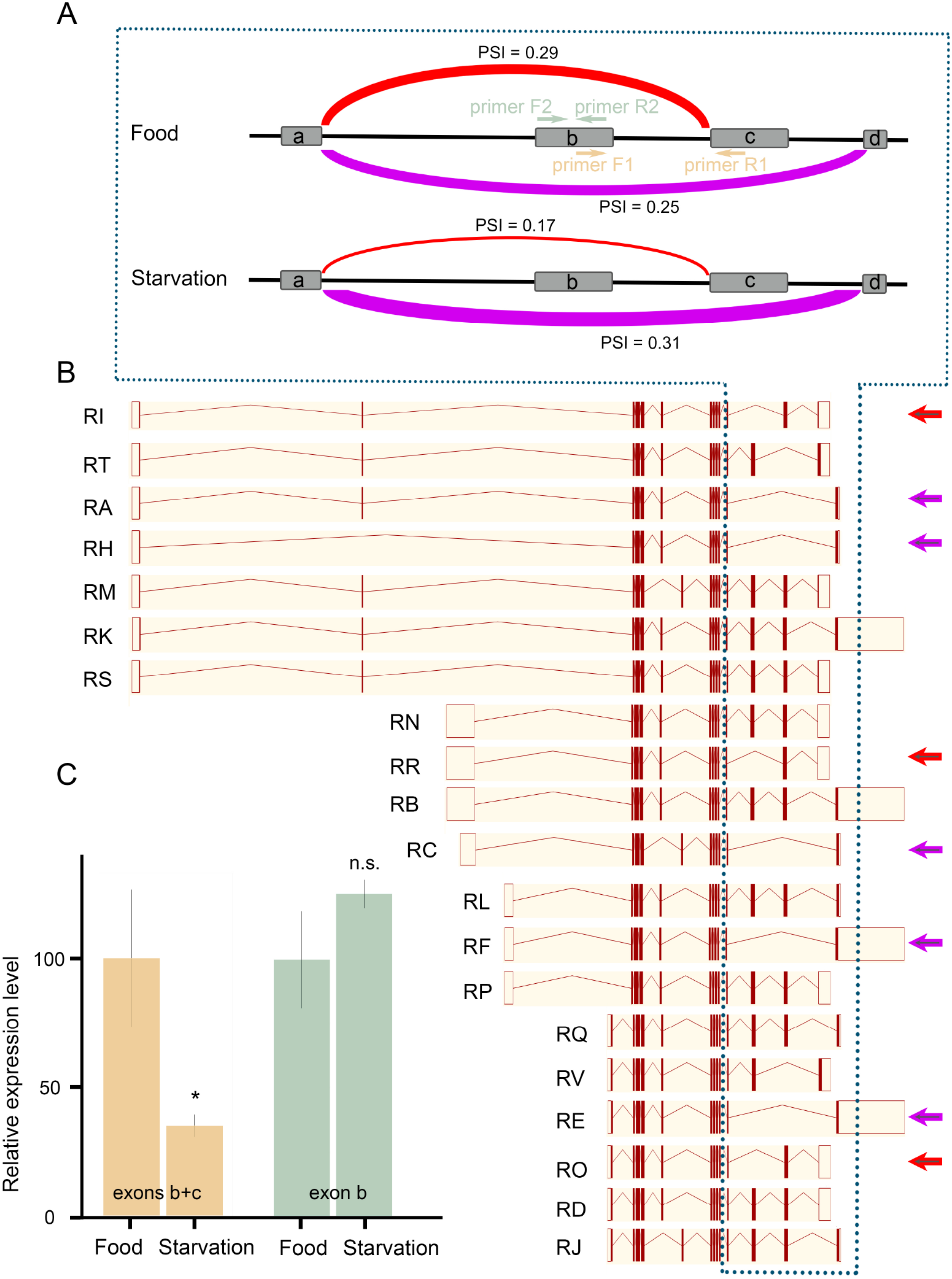
*syp* is alternatively spliced under starvation. (A) Differential usage of exons in *syp* under food and starvation conditions. Four exons involved in the alternative splicing were shown and referred to as exon a, exon b, exon c, and exon d. PSI: Percentage Sliced In. (B) Various splice variants of *syp* (www.flybase.org). Arrows indicate splice variants that were either downregulated (red color) or upregulated (purple) under starvation. (C) Relative expression of targeted exons measured by qPCR. The targeted locations of each pair of primers were indicated in panel A. * *P* < 0.05, n.s.: *P* > 0.05, unpaired *t*-test, n = 4 per condition. Error bars represent SEM.

To identify genes that underwent alternative splicing under starvation, we analyzed RNA-seq data using LeafCutter, a software that can identify and quantify both novel and known alternative splicing events [33]. LeafCutter focuses on intron excisions and groups RNA-seq reads into different clusters based on their mapped location to the genome. Therefore, the final data presentation is in the form of clusters that include various splicing forms. A cluster was considered significant when FDR < 0.05. A total of 1927 clusters from 359 genes were alternatively spliced under starvation (S7 Table). Gene enrichment analyses showed that these genes were implicated in a variety of biological processes, including regulation of mRNA splicing itself (S8 Table). Remarkably, the most significant molecular function GO term is mRNA binding (S8 Fig, S8 Table), suggesting that upon starvation, genes encode RNA-binding proteins tend to be alternatively spliced.

Among these 1927 significant clusters, 359 of them, from 180 genes, have a ΔPSI larger than 10%. PSI, Percentage Sliced In, is the fraction of a gene’s mRNAs that contain the exon; therefore, ΔPSI is the difference in PSI between the starvation condition and the food condition. Among these 359 clusters, 80.8% of them have small splicing changes (10% < ΔPSI < 25%), 18.7% have intermediate splicing changes (25% ≤ ΔPSI ≤ 50%), and 0.5% have large splicing changes (ΔPSI > 50%). Gene *syp* is alternatively spliced under starvation (S7 Table). A total of 10 splicing events were found in this significant cluster. Among them, the top two ΔPSIs were −12% and 6% (Fig 5A). The splice variants that correspond to the changes are *syp*-RI (Transcript ID: FBtr0334711), *syp*-RR (FBtr0334720), *syp*-RO (FBtr0334717), *syp*-RA (FBtr0083958), *syp*-RH (FBtr0113256), *syp*-RC (FBtr0083960), *syp*-RF (FBtr0083961), and *syp*-RE (FBtr0083963) (Fig 5B).

To confirm that these changes were not from sequencing noises, we designed PCR primers to target the middle exons within this region (exons b and c, Fig 5A) and performed RT-qPCR. Results showed that the expression of exons b and c in starved flies was reduced to 38% of that in flies maintained under the food condition (Fig 5C), which confirmed the increased splicing from exon a to exon d. We further designed primers to target exon b only and found that the expression of exon b alone was unchanged after starvation (Fig 5C), which was not surprising since this exon was involved in two splicing events that were regulated in opposite directions by starvation. Therefore, *syp* was alternatively spliced upon starvation. It is likely that specific *syp* splice variants play the role in regulating SIH.

### Knocking out two alternatively spliced exons in *syp* leads to reduced SIH

To study the significance of alternative splicing involving exons b and c in SIH, we generated exons b and c deletion flies using CRISPR/Cas9 [34, 35]. We designed sgRNAs to target both exons (Fig 6A). Complete deletion of exons b and c was confirmed in the mutant allele *syp*^1^ by sequencing. The *syp*^1^ allele was initially maintained over *TM3, Ser*. We observed very few homozygous mutant flies emerged from the breeding vials (4 homozygotes out of 457 flies), suggesting that deletion of these two exons potentially affects either development or adult viability.

**Fig 6.**
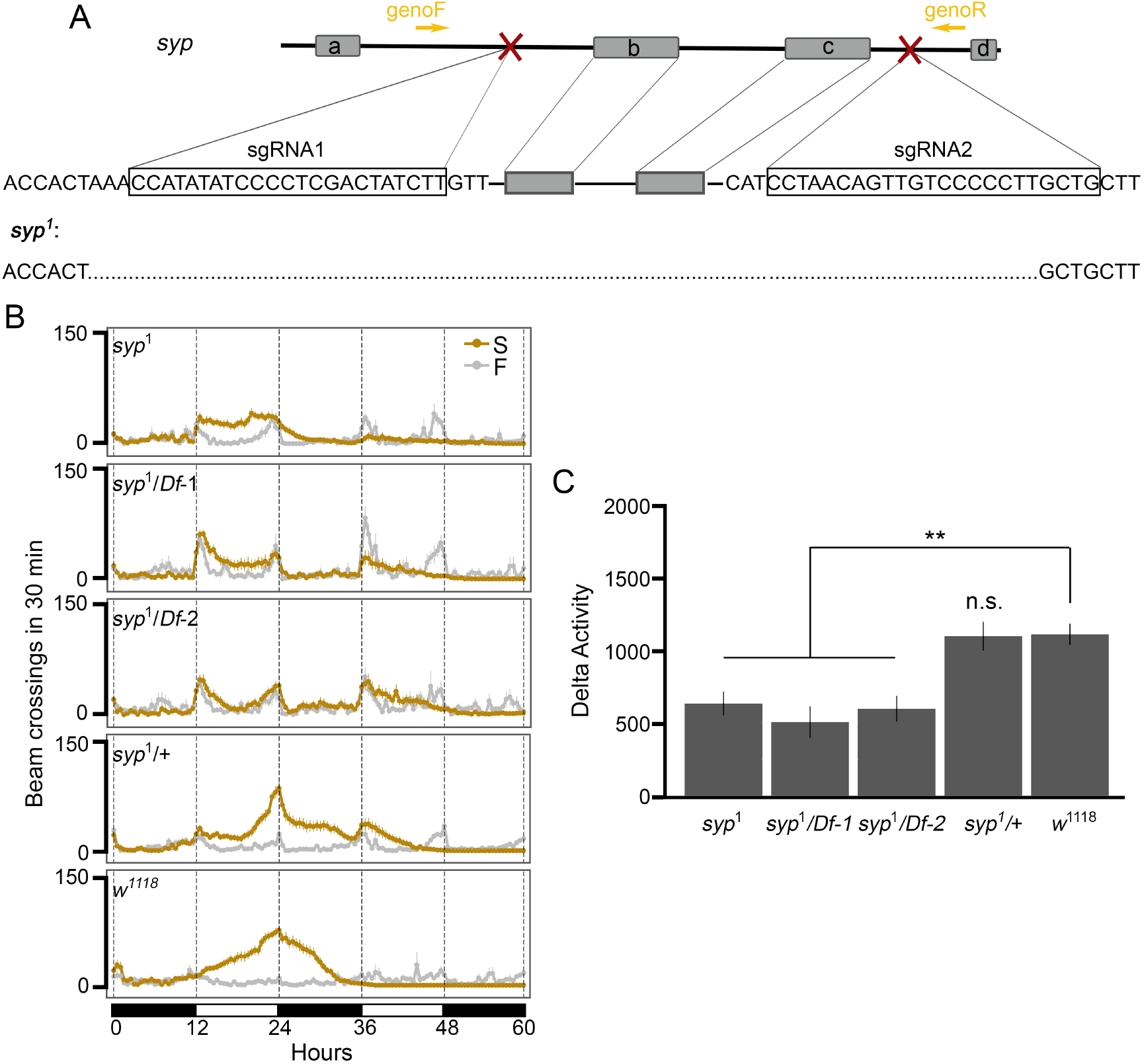
Deletion of exons b and c in *syp* leads to reduced SIH. (A) Generation of *syp* exons b and c deletion lines using CRISPR/Cas9. Two sgRNAs that target the intronic regions adjacent to exon b and exon c (symbol X in red) cut each side of the targeted region, respectively. The sequences of sgRNAs were show in the rectangles. A pair of primers (genoF and genoR) were used to amplify the targeted region for sequencing. Exons b and c were deleted in the *syp*^1^ allele based on sequencing results. (B) Locomotor activity responses to starvation from each genotype under either food (F) or starvation condition (S). n = 16 - 26 per genotype per condition. *Df-1*: *Df(3R)BSC124*; *Df-2*: *Df(3R)BSC141*. (C) Summary plot of SIH from each genotype in panel B. ** *P* < 0.01, ** *P* < 0.001, n.s.: *P* > 0.05, unpaired *t*-test with Bonferroni Correction. Error bars represent SEM.

To determine the stage of the lethality occurred, we outcrossed *syp*^1^ to *w*^1118^;*TM3* /*TM6B, Hu, Tb* and maintained the *syp*^1^ allele over *TM6B, Hu, Tb* so that we could study pupae by taking advantage of the phenotypic marker Tubby (Tb) associated with *TM6B, Hu, Tb*. We observed a normal ratio between non-Tb and Tb pupae (140 non-Tb vs. 291 Tb, Chi-squared test for given probabilities, *P* = 0.7223), demonstrating that exons b and c are dispensable for development before pupal stages. We also noticed that after outcrossing, although the number of the *syp*^1^ homozygous flies was still lower than expected (9 homozygotes out of 194 flies, that is ∼5%), but higher than the initial observation (∼1%). The requirement of *syp* to the adult survival is consistent with a previous study showing that flies from two *syp* mutant lines, *syp*^f03775^ and *syp*^e00286^, have decreased adult viability, but mutants can survive until late pupal stages [28]. Notably, insertion sites in *syp*^f03775^ and *syp*^e00286^ are within the C-terminal region of *syp*. Specifically, f03775 is located in the intronic region between the two exons deleted in our deletion lines and e00286 is located immediately upstream of these two exons.

Next, we examined the locomotor responses of survived *syp*^1^ homozygous flies to starvation. Results from locomotor activities under either food or starvation conditions showed that compared to *w*^1118^ flies, *syp*^1^ homozygotes had significantly reduced SIH (*P* = 1.11e-4, Fig 6B and 6C), similar to neuron-specific knockdown flies (Fig 3). We also examined *syp*^1^ heterozygous flies and did not see a difference in SIH between these flies and *w*^1118^ (*P* = 0.9162).

To further confirm that the reduced SIH in *syp*^1^ homozygotes was indeed due to the deletion of exons b and c in *syp*, we generated and tested *syp*^1^/*Df(3R)BSC124* and *syp*^1^/*Df(3R)BSC141* flies. *Df(3R)BSC124* and *Df(3R)BSC141* are two independent deficiency alleles in which a number of genes including *syp* are deleted (www.flybase.org). Therefore, *syp*^1^/*Df(3R)BSC124* and *syp*^1^/*Df(3R)BSC141* represent a homozygous deletion for exons b and c on an otherwise heterozygous *syp* background. Similar to *syp*^1^ homozygotes, flies from both *syp*^1^/*Df(3R)BSC124* and *syp*^1^/*Df(3R)BSC141* also showed reduced SIH (*P* = 8.55e-05 and *P* = 8.18e-05, respectively). Together, these results demonstrate that alternative splicing of *syp* involving exons b and c play an essential role in regulating SIH in adult flies.

## Discussion

Starvation-Induced Hyperactivity (SIH) has been observed in different species [3–12], suggesting that it has a genetic component. SIH facilitates food acquisition and energy intake when food is available [7], but it increases energy expenditure and makes starved animals even more vulnerable when food is not available [4]. Therefore, genetic dispositions to either too much or too little SIH would impair fitness depending on the environment. Taking advantage of the genetic diversity in the DGRP strains derived from a wild population, we have demonstrated for the first time the significant genetic contribution to SIH. Our studies show that the broad sense heritability (*H*^2^) of SIH is 0.38, which is lower than the reported heritability of developmental traits in the DGRP (e.g. *H*^2^ is 0.89 for developmental time [36], 0.66-0.88 for pigmentation [37], and 0.71-0.78 wing morphology [38]), but is higher than the reported heritability of behavioral traits in the DGRP (e.g. *H*^2^ is 0.03-0.09 for courtship behavior [39] and 0.02-0.45 for olfactory behavior [40–42]), suggesting that SIH has a relatively strong genetic basis compared to other behavioral traits.

Our genome-wide association studies identified 23 loci from 14 genes significantly associated with SIH in adult *Drosophila* (Fig 2A, Table 1, Table 2). Gene enrichment analyses revealed that mRNA binding proteins were important for SIH (Fig 2C). We validated the role of RNA-binding protein Syp in adult neurons in regulating SIH (Fig 3, Fig 4) and showed that specific *syp* splice variants were responsible for SIH (Fig 5, Fig 6). Although the present study is the first unbiased genome-wide study on SIH, other candidate gene approaches have discovered a number of SIH genes in *Drosophila* in recent years, which includes *AMP-activated protein kinase* (*AMPK*) [5, 6], *Adipokinetic hormone* (*Akh*) [4], *Adipokinetic hormone receptor* (*AkhR*) [8], *Insulin-like peptides* (*Ilps*) [8], *Insulin-like receptor* (*InR*) [8], *Tyramine beta hydroxylase* (*Tbh*) [7], and *dG9a* [9]. Majority of these candidates play important roles in relaying metabolic information to neurons. Expression of some of these genes were regulated by starvation, as shown in our differential expression analyses from RNA-seq data. For example, the mRNA levels of both *AkhR* and *InR* were significantly altered after starvation (logFC = −1.08 and logFC = 1.45; *P* = 1.65e-9, and *P* = 3.15e-23 for *AkhR* and *InR*, respectively. S6 Table). The present study further suggests that RBPs and post-transcriptional regulation could be another important pathway that regulates SIH. Moreover, our RNA-seq data show that only 15 genes (3 up-regulated and 12 down-regulated) are shared between the 658 differentially expressed genes and the 180 alternatively spliced genes, suggesting that regulation at both transcriptional and post-transcriptional levels for the same gene is uncommon in response to starvation, at least in the head. Gene enrichment analyses further suggest that metabolic genes tend to be regulated at the transcriptional level while genes involved in the mRNA regulation tend to be regulated post-transcriptionally (S6 Fig). This is also shown in our splice variants analysis followed by GO analysis. Genes encoding RBPs seem to be especially regulated by alternative splicing (S8 Fig).

*syp* has a number of splice variants (Fig 5B). These variants mainly differ at the N-terminus and the C-terminus, whereas the remaining middle region of the protein, corresponding to the three RNA recognition motifs, are largely unchanged. The sequence discrepancy among various splice variants implies a potential role of UTRs in response to different stimuli. Our results from RNA-seq demonstrate that under starvation, *syp* is mainly regulated at the C-terminus (Fig 5A). We postulate that either Syp itself or some other RBPs may regulate the splicing of *syp* under starvation. Such self-regulatory or cross-regulatory networks in the alternative splicing of RBPs have been reported in previous studies [43]. Moreover, alternatively spliced exons in *syp* seem not only involved in regulating starvation responses. We have found that deletion of these axons leads to semi-lethality in adult flies. Although the argument is beyond the scope of the current study, our KO data suggest that the SIH and lethality might be mediated through different splice variants. Given that the skipping of exons b and c is elevated upon starvation (Fig 5A) and that knockout of them leads to reduced SIH (Fig 6B), it is very likely that transcripts with no exons b and c are responsible for Syp’s role in SIH. In comparison, the survival phenotype is more likely due to the absence of the transcripts containing either exon b or c or both.

Syp is a *Drosophila* homolog of human SYNaptotagmin-binding Cytoplasmic RNA-Interacting Protein (SYNCRIP)/hnRNP Q, and they share 47% sequence identity [28]. Similar to the mammalian SYNCRIP, Syp consists of three RNA recognition motifs and one acidic domain at the N-terminus [28, 44]. However, it lacks the arginine-glycine-glycine domain at the C-terminus [28]. In mammals, different domains mediate the interaction of SYNCRIP with different effectors, which renders SYNCRIP a wide range of functions including circadian regulation [45–47], neuronal morphogenesis [48–50], and stress response [51]. Misregulation of SYNCRIP has been reported in neurodegenerative diseases [52, 53], psychiatric disorders [54, 55], and cancer [56, 57]. In flies, studies have shown that Syp is required for different developmental phenotypes, such as oogenesis [28], maintaining a normal structure and function of synapses at the neuromuscular junction in larvae [29, 30], and determining neuron/glia cell fates [31, 32]. The present finding is the first indication that Syp has a unique function in adult flies (Fig 3) independent of its developmental effects (Fig 4). At the subcellular and molecular level, previous studies have shown that Syp, in muscle cells at the *Drosophila* larval neuromuscular junction, can modulate the presynaptic vesicle release through regulating postsynaptic translation of a retrograde signal [30] and can regulate activity-dependent synaptic plasticity [58]. Similar functions have also been observed in mammalian SYNCRIP such that it is a component of neuronal RNA transport granules that can regulate dendritic morphology [48, 59–61], and that it inhibits translation by competing with the poly(A) binding protein [62]. We hypothesize that Syp might be able to quickly affect protein synthesis in neurons and even local protein synthesis in synapses in response to starvation. Further studies are needed to examine the molecular mechanisms, including the upstream and downstream pathways associated with Syp in adult neurons in response to starvation.

In summary, we report genome-wide association studies of SIH as a behavioral trait. Our data suggest a strong genetic component in SIH. We further report an essential role of *syp* and post-transcriptional regulation in adult neurons in regulating SIH.

## Materials and methods

### Drosophila stocks

The *Drosophila melanogaster* Genetic Reference Panel (DGRP) strains were from the Bloomington Drosophila Stock Center (BDSC). The UAS-*syp* RNAi lines (#33011, #33012) and the genetic control line were from the Vienna Drosophila Resource Center (VDRC). The *actin*-Gal4 (#4414), the pan-neuronal driver *nSyb*-Gal4 (#51635), *Df(3R)BSC124* (#9289), and *Df(3R)BSC141* (#9501) were from the BDSC. Unless stated otherwise, all flies were reared on the standard cornmeal medium from FlyKitchen at the University of Chicago at 25 °C under a 12-hr:12-hr light: dark cycle with the light on at 06:00 and off at 18:00.

### Locomotor assay setup

Male flies, 1 to 3 d old, were anesthetized briefly and transferred into activity tubes filled with the standard cornmeal medium where they were allowed to recover from CO2. One day later, flies were randomly separated into two groups. Flies in the first group were transferred into activity tubes filled with 4% sucrose plus 5% yeast in 1% agar (food condition), and flies in the second group were transferred into activity tubes filled with 1% agar only (starvation condition). The other end of activity tubes was inserted with a small cotton ball to allow air exchange for the fly and also to prevent fly from escaping. The locomotor activity of each fly was monitored using the Drosophila Activity Monitor (DAM2, Trikinetics Inc.). Unless stated otherwise, the assay chamber was maintained at 25 °C under a 12-hr:12-hr light: dark cycle with the light on at 06:00 and off at 18:00. The start point on the plots is the light off point, which is 18:00 on the setup day, which is about 6-7 hours after flies were transferred to the activity tubes. We used 8 flies per condition per genotype for screening. The number of flies in other experiments was indicated in the text.

### Data analysis of the locomotor responses to starvation

Infrared beam breaks per 30-minute interval from two conditions were averaged and plotted as a function of time point for each strain. The total activity of each fly from a 12-hour period (either daytime or nighttime) in each condition was first summed, and the activity difference between the two conditions was then calculated. Since flies were monitored for a total of sixty hours, there were five DAs. The largest DA among five DAs was referred to as DA of each fly. The averaged DA from 8 flies was used to represent the strain DA.

### Starvation Resistance

Starvation Resistance is represented by the time point (0-120) when all eight flies under starvation were dead. The death of a fly was inferred from the activity data such that there were no beam crosses. In the case of all flies survived to the last time point, an index of 121 was used.

### Genome-wide association analysis

Genome-wide association analysis on SIH was performed using the DGRP analysis pipeline at http://dgrp2.gnets.ncsu.edu [16, 17]. Briefly, each raw dataset was first adjusted for the Wolbachia infection and the inversion with a linear model where the raw dataset as the response variable and the infection status and five major inversion polymorphisms as covariates (S3 Table). Residuals from this linear model were then used as response variable to fit a mixed linear model: Y = *µ* + G + r, where *µ* is the overall population mean for each trait, G is the effect of SNPs or INDELs being tested, and r is a polygenic component with covariance between lines determined by their genome relationship [17]. A total of 1,905,103 SNPs or INDELS with minor allele frequencies (MAFs) larger than 0.05 were included in the analysis. A nominal *P*-value threshold of *P* < 1 × 10^−5^ was used for declaring SNPs or INDELs to be significantly associated with trait variation by following previous studies [16, 17].

### Gene enrichment analysis

The web-based enrichment analysis tool FlyEnrichr [24, 25] was applied to evaluate the list of candidate genes, and to obtain a set of enriched functional annotations in three domains: biological process (BP), cellular component (CC), and molecular function (MF) [63] or in KEGG (Kyoto Encyclopedia of Genes and Genomes) pathways [64]. GO or KEGG terms with *P* < 0.05 (after Benjamini-Hochberg correlation) were listed in S5 Table and S8 Table.

### Western blotting

For the Gal80^ts^ experiment, a group of 25-30 male flies, 1 to 3 d old, were reared on the standard cornmeal medium at either 18°C or 31°C for two days. For the protein level of Syp in *w*^1118^ flies under either food or starvation conditions, a group of 25-30 male flies, 1 to 3 d old, were maintained in either food or starvation conditions at the room temperature for one day. The total protein from adult heads was then extracted and quantified. A total of 50 *µ*g protein from each sample was loaded for SDS-PAGE. Separated proteins were electrophoretically transferred to the PVDF membrane. After blocking, the membrane was incubated with primary antibodies (Guinea pig anti-Syp 1:2000 Ilan Davis, UK; Mouse anti-tubulin 1:500 Developmental Studies Hybridoma Bank, DSHB) and then the secondary antibodies (peroxidase-labeled anti-Guinea pig IgG 1:10,000, Sigma; peroxidase-labeled anti-mouse IgG 1:10,000, Jackson ImmunoResearch Laboratories). Signals were detected with enhanced chemiluminescence (ThermoScientific).

### RNA extraction

A group of 30 male flies, 1 to 3 d old, were maintained under either food or starvation conditions for one day. Four biological samples were collected for each condition. Total RNA was extracted from heads using RNA extraction kit (Zymo Research). After the removal of genomic DNA (DNA-free kit, AMBION), total RNA was quantified using a NanoDrop spectrophotometer (Thermo Scientific), and the concentrations of total RNA were then diluted to 100 ng/*µ*L.

### RNA sequencing and data analysis

RNA sequencing was performed in the Genomics Core Facility at the University of Chicago. Briefly, the integrity of total RNA was assessed using a bioanalyzer 2100 (Agilent), and poly(A)+ RNA-seq libraries were then prepared from each sample, multiplexed, and sequenced by 100-bp paired ends using Illumina HiSeq4000 sequencer (S3 Fig, S4 Fig). Sequencing was duplicated using two flow cells. A total of 15-30 million reads were generated for each library per sequencing batch. Sequencing quality was assessed using FastQC [65], and the mean quality score ranges from 38.84 to 39. Raw sequencing reads from each group (treatments + biological replicates + sequencing batches) were individually mapped to the Drosophila genome (v6.22) using STAR (v2.6.1b) [66]. About 95% of raw reads from each sample were uniquely mapped to the genome. Each bam file was then processed using ASpli [67] to obtain differentially expressed genes and thereby to access the sequencing batch effect (S5 Fig). Since results from two sequencing batches are highly correlated (*ρ* = 0.99, *P* < 2.2e-16, Pearson correlation test), we merged two sequencing bam files from the same sample using samtools [68] for downstream analyses. Alternative splice variants were analyzed using Leafcutter [33].

### RT-qPCR

The cDNAs were synthesized using SMARTScribe Reverse Transcription Kit (TaKaRa). Two pairs of primers that target either both exons b and c or exon b only were used to amplify these two exons. Primer sequences of the first pair of primers are: 5’-TTC ACC GAT GGC TAG TGG AC and 5’-GTT GGC CAA CGA CTC TGC CA, and primer sequences of the second pair of primers are: 5’– TTC GGT TTC TCG GAC TAT CG and 5’ – CCA CCG TTC GGG TAA TCA TA. One pair of primers that targeted the house-keeping gene *ribosomal protein 49* (*rp49*) was used as the internal control for qPCR. Primer sequences are: 5’-GCT AAG CTG TCG CAC AAA TG and 5’-GTT CGA TCC GTA ACC GAT GT.

### Generation of exon b and exon c deletion using CRISPR/Cas9

The deletion allele was generated using CRISPR/Cas9 [34,35]. The sgRNAs were designed using FlyCRISPR Optimal Target Finder (http://targetfinder.flycrispr.neuro.brown.edu/) and *in vitro* transcribed using T7 transcript kit (MEGAscript T7 Transcription Kit, Cat # AMB13345). The sequences of sgRNAs were shown in Fig 6A. The sgRNAs and Cas9 protein (RNA bio company, Cat #CP01) were injected into embryos from flies with a genotype: *nos-phiC31, y[1] sc[1] v[1] sev[21]; Py[+t7*.*7]=CaryPattP2* (BDSC #25710). Deletion alleles were maintained over *TM3, Ser* (BDSC #4534). Deletion was confirmed by sequencing the targeted region amplified using a pair of primers (Fig 6A). Primer sequences are 5’-CGA ACT CTC TGT GTC GCA AG (genoF) and 5’-AGT TGG CAT TGG ATT GGT GT (genoR).

### Correlation, quantitative genetic, and statistical analyses

All analyses were performed using the statistical software R v3.6.1 (http://www.r-project.org). Pearson correlation test was applied for correlation analyses. Other statistical tests were indicated in the text or figure legends. The broad-sense heritability (*H*^2^) of SIH was computed as *H*^2^ = *σ*^2^_G_/(*σ*^2^_G_ + *σ*^2^_E_), where *σ*^2^_G_ is the among-line variance component and *σ*^2^_E_ is the error variance.

## Data availability

All raw and processed RNA-sequencing data generated in this study have been submitted to the NCBI Gene Expression Omnibus (GEO) under accession number GSE149790. The complete GWAS dataset has been deposited to zenodo.org (doi: https://doi.org/10.5281/zenodo.4091399).

## Supporting information

**S1 Fig. Setup image of the locomotor activity assay**. One end of the activity tube (Diameter × Length = 5 mm x 65 mm, Trikinetics Inc.) was filled with either 4% sucrose plus 5% yeast in 1% agar in the food condition or 1% agar only in the starvation condition. The length of the medium is 1 cm. The other end of the tube was stopped with a small cotton ball. One male fly was introduced into each tube.

**S2 Fig. Linkage Disequilibrium map and Quantile-Quantile plot for GWA on SIH**.

**S3 Fig. Experimental design and workflow of RNA sequencing. S4 Fig. Integrity of total RNA from each sample**.

**S5 Fig. Correlation analyses of genes from two sequencing batches**. A: Scatter plot for all genes. A total of 9452 genes were included in the analysis. Each dot represents one gene. Dots with cyan color are the genes that show the same direction of change after starvation for two batches. Dots with yellow color are the genes that show the opposite direction of change after starvation for two batches. B: Scatter plot for genes with false discovery rate (FDR) < 0.05 and fold change ≥ 2 or fold change ≤ −2.

**S6 Fig. Differential expression analysis**. A: Clustering and heatmap showing mRNA levels in four biological replicates of each condition. B: Volcano plot of significantly regulated genes by starvation (Cutoff values: FDR < 0.05 and fold change ≥ 2 or fold change ≤ −2). (C) Enriched KEGG pathways associated with either down-regulated (blue bars) or up-regulated (red bars) genes.

**S7 Fig. Western blot and quantification of Syp protein level**. A: Western blot of adult fly head homogenate from *w*^1118^ flies under either food or starvation conditions. Four biological replicates per condition. Tubulin is the loading control. B: Quantification of Syp from panel A. n.s: *P* > 0.05, unpaired *t*-test. Error bars represent SEM.

**S8 Fig. Molecular Function GO terms associated with alternatively spliced genes**. Only terms associated with more than four genes were shown in this plot. See the complete list in S8 Table.

**S1 Table. Data of SIH and of Starvation Resistance for DGRP strains. S2 Table. Analysis of variance of SIH in the DGRP**.

**S3 Table. Covariates analyses**.

**S4 Table. Gene lists of GWA with *P*** < **1** × **10**^**-4**^, ***P*** < **8** × **10**^**-5**^, ***P*** < **6** × **10**^**-5**^, ***P*** < **4** × **10**^**-5**^, **and *P*** < **2** × **10**^**-5**^.

**S5 Table. Over-representation of Gene Ontology Categories associated with nominated genes with *P*** < **1** × **10**^**-4**^ **or *P*** < **8** × **10**^**-5**^.

**S6 Table. Differentially expressed genes. S7 Table. Alternatively spliced genes**.

**S8 Table. Over-representation of Gene Ontology Categories and KEGG pathways associated with alternatively spliced genes**.

## Acknowledgments

We thank Ilan Davis for sharing the anti-Syp antibody. We thank Chunyu Liu and Xiaochang Zhang for helpful discussions. We thank Yang I Li, Jack Humphrey, Julian Beckman, Cai Qi, Shuaibo Han, Wei A Du, and Meredith Wells for technical assistance. We thank the Bloomington Drosophila Stock Center (BDSC) and the Vienna Drosophila Resource Center (VDRC) for fly stocks. The computational analysis in this study was performed with resources provided by the University of Chicago Research Computing Center. This work was supported by the National Institutes of Health (T32MH020065 & T32DA434693 to W.C., R01GM100768 to X.Z.).

