## Supplementary figures and images for "RNA-binding protein Syncrip regulates Starvation-Induced Hyperactivity in adult *Drosophila*"

### S1 Fig

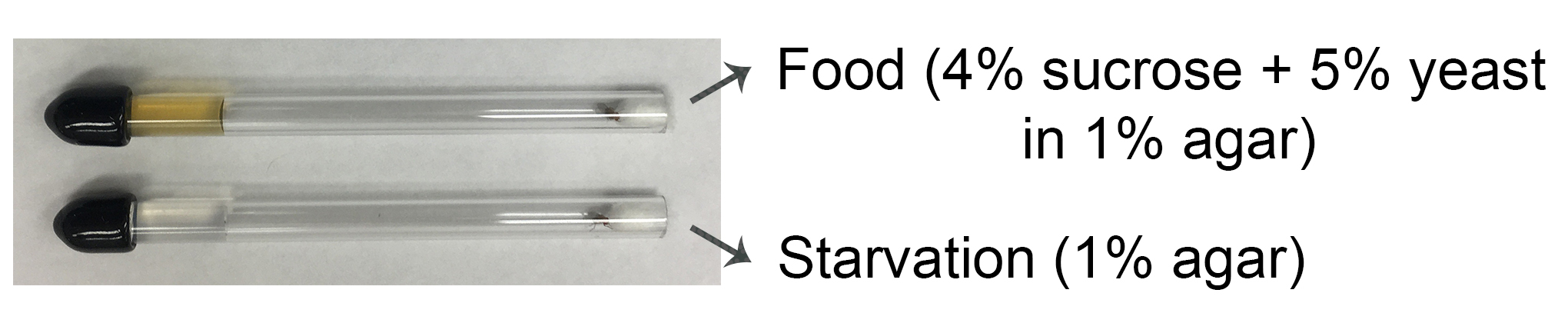

### S2 Fig

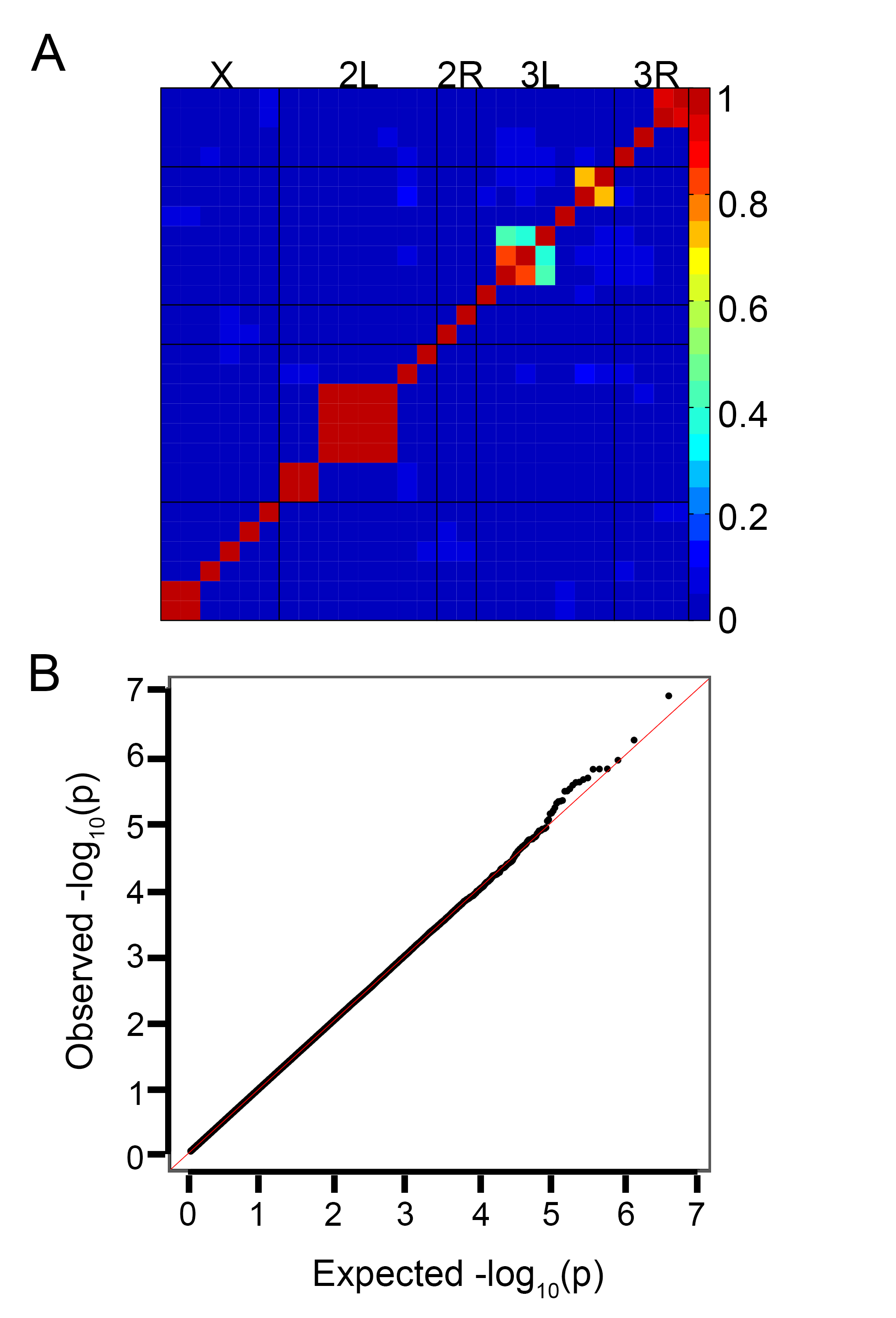

### S3 Fig

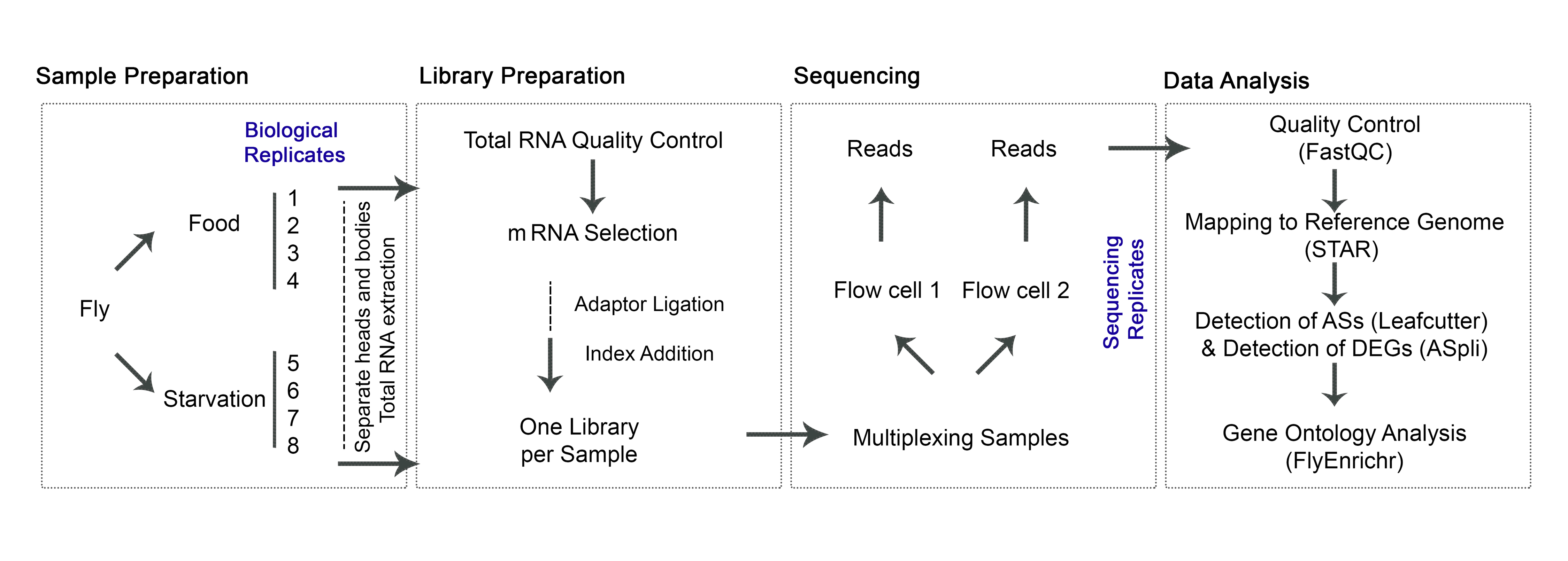

### S4 Fig

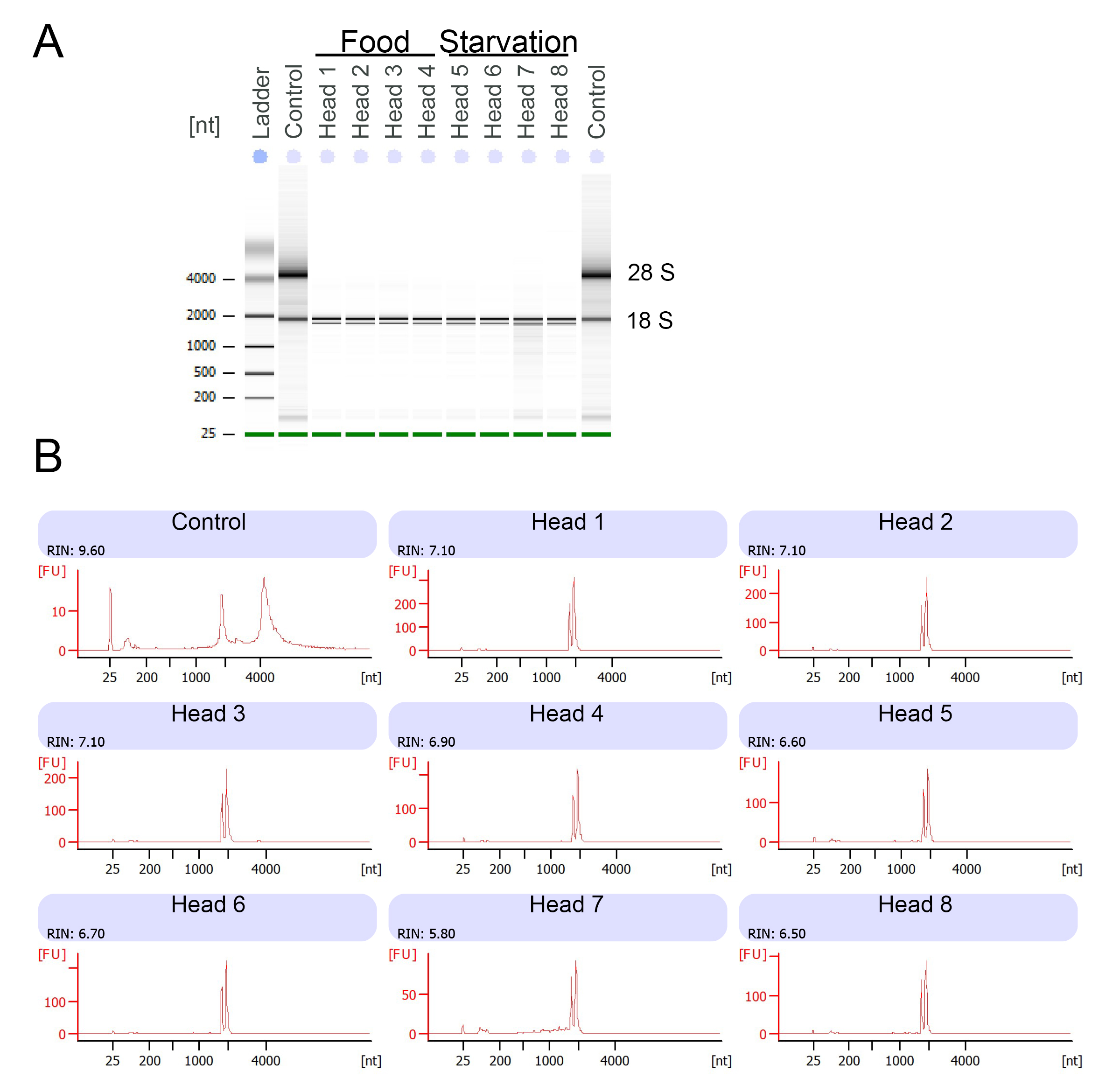

### S5 Fig

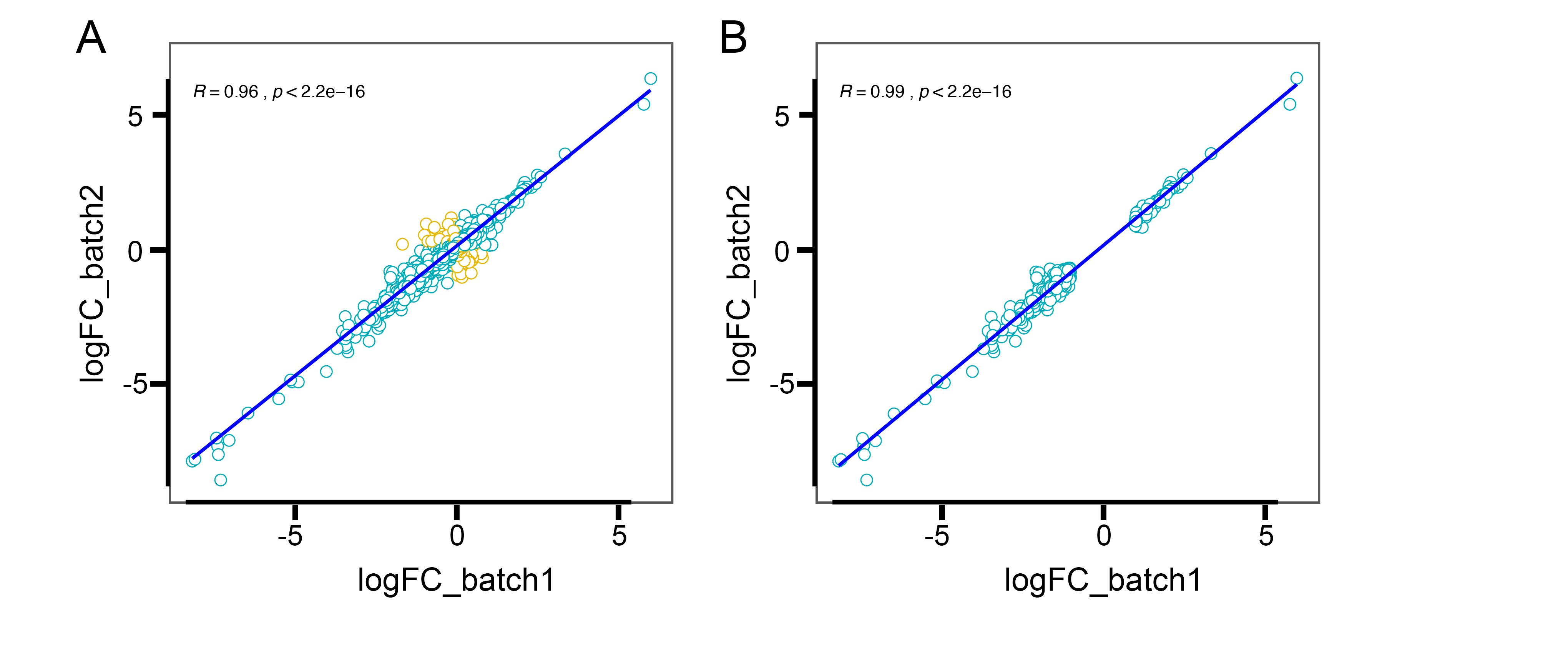

### S6 Fig

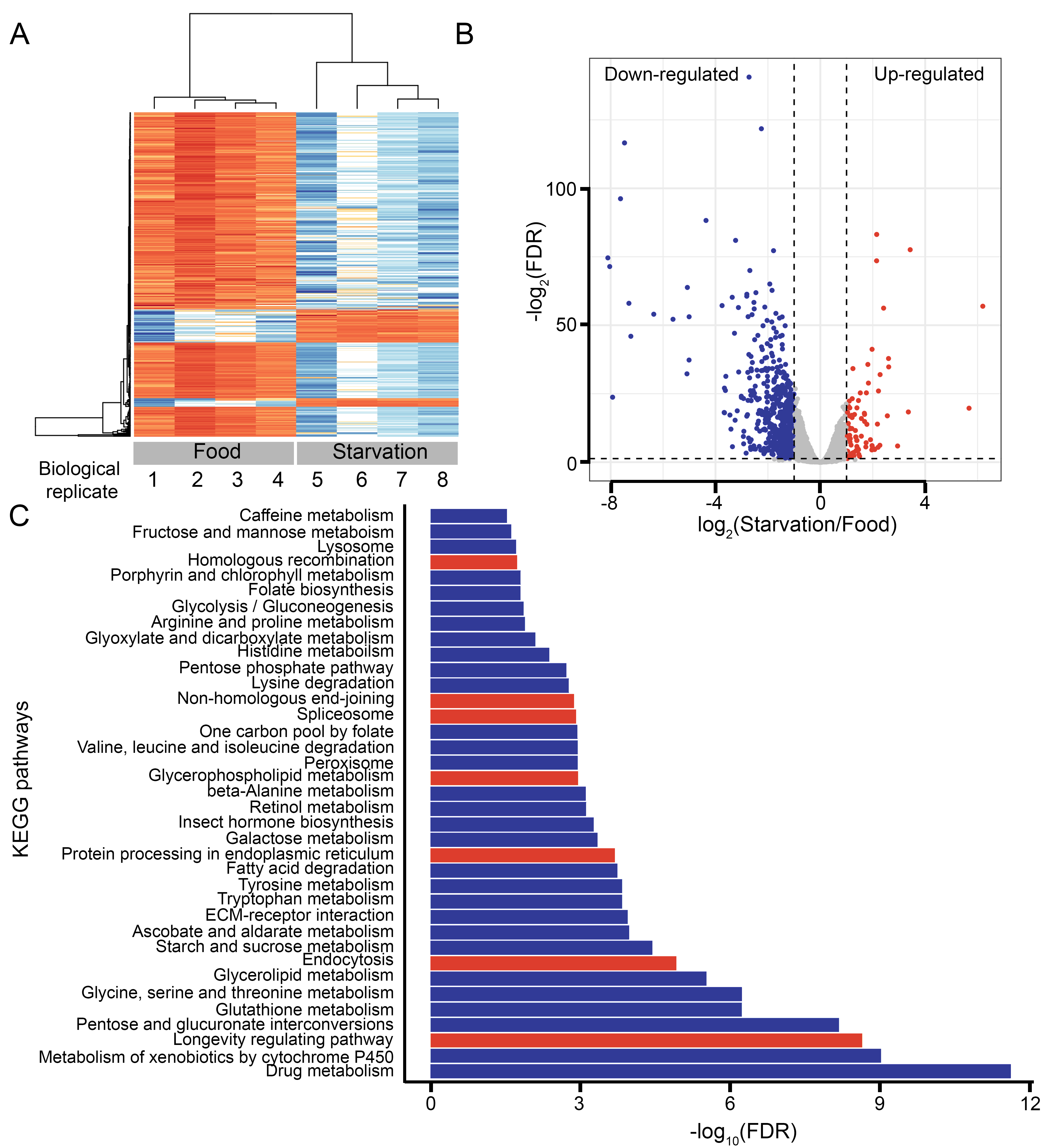

### S7 Fig

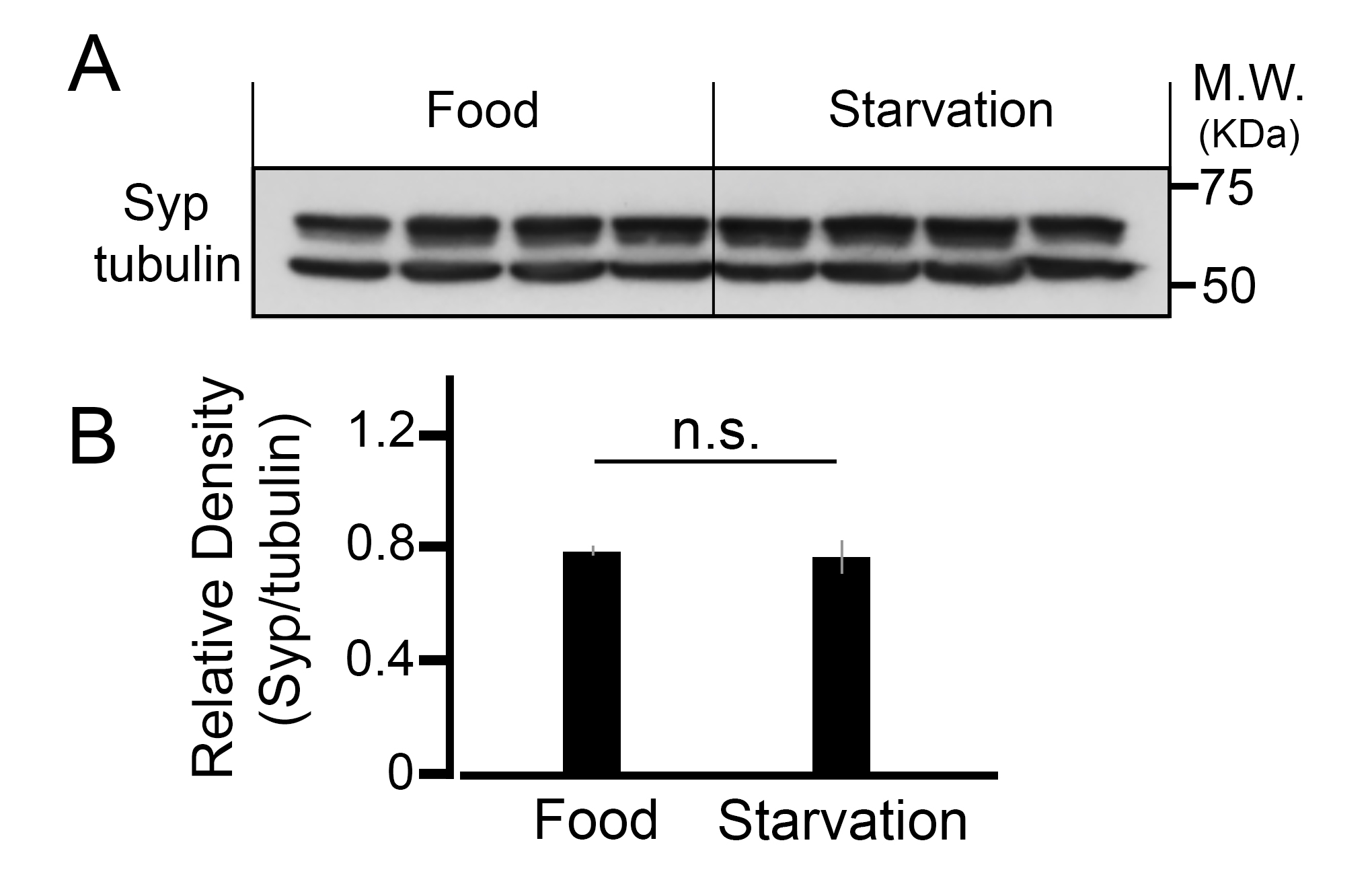

### S8 Fig

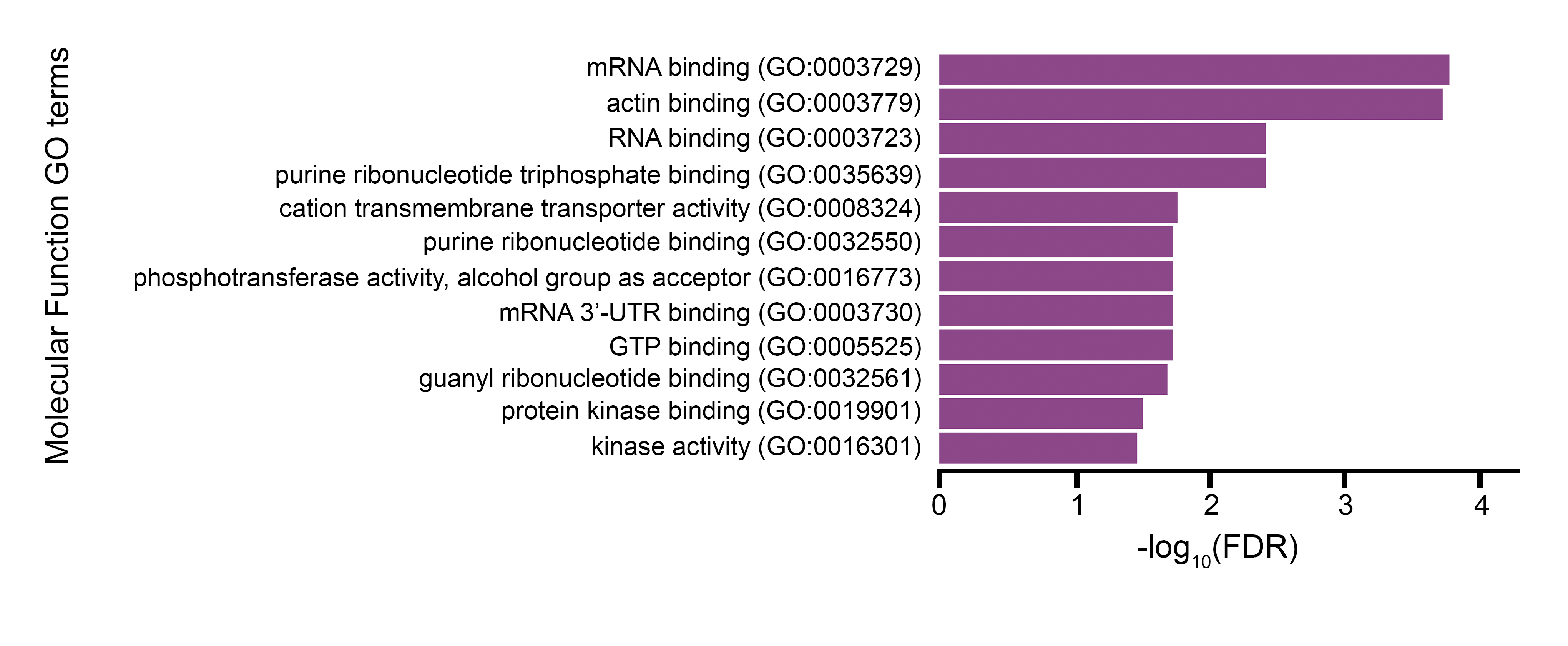
